# Stereomotion processing in the non-human primate brain

**DOI:** 10.1101/638155

**Authors:** Yseult Héjja-Brichard, Samy Rima, Emilie Rapha, Jean-Baptiste Durand, Benoit R. Cottereau

**Author notes:** Corresponding authors: Yseult Héjja-Brichard & Benoit R Cottereau.

## Abstract

The cortical areas that process disparity-defined motion-in-depth (i.e. cyclopean stereomotion) were characterised with functional magnetic resonance imaging in two awake, behaving macaques. The experimental protocol was similar to previous human neuroimaging studies. We contrasted the responses to dynamic random-dot patterns that continuously changed their binocular disparity over time with those to a control condition that shared the same properties, except that the temporal frames were shuffled. A whole-brain voxel-wise analysis revealed that in all four cortical hemispheres, three areas showed consistent sensitivity to cyclopean stereomotion. Two of them were localised respectively in the lower bank of the superior temporal sulcus (CSM_STS_) and on the neighbouring infero-temporal gyrus (CSM_ITG_). The third area was situated in the posterior parietal cortex (CSM_PPC_). Additional ROIs-based analyses within retinotopic areas defined in both animals indicated weaker but significant responses to cyclopean stereomotion within the MT cluster (most notably in areas MSTv and FST). Altogether, our results are in agreement with previous findings in both human and macaque and suggest that the cortical networks that process cyclopean stereomotion is relatively well preserved between the two primate species.

## Introduction

Motion perception is a fundamental property of the visual system in most animal species. It enables to track over time the position of elements in a scene and thereby facilitates navigation or interactions with moving objects. Historically, numerous studies have characterised planar (i.e. 2D) motion processing in the primate nervous system (see e.g. Maunsell and Newsome, 1987). In macaque, single-cell recordings showed that it is computed at the cortical level within an extended network that begins in the primary visual cortex and includes higher-level visual areas such as area V4 in the ventral patway (Li et al., 2013) and area V3A (Galetti et al., 1990) or V6 (Pitzalis et am., 2013) in the dorsal pathway. Within the Superior Temporal Sulcus (STS), area MT notably hosts neurons whose responses are highly selective to motion direction (see e.g. Maunsell & Newsome, 1987) and also reflects motion perception (Newsome & Paré, 1988; Britten et al., 1996). In human, neuroimaging studies suggested that 2D motion is also processed within an extended network that includes a putative homologue of area MT: hMT (Huk et al., 2002). Over the last 20 years, the emergence of monkey fMRI (see e.g. Vanduffel et al., 2001) made possible the further characterisation of the correspondence between the cortical areas involved in motion processing in the two species. This comparative approach revealed that 2D motion engaged largely similar networks in macaques and in humans. It notably suggested that motion processing in MT and its satellite areas (V4t, FST, and MSTv) on the one hand and in hMT and its satellites (pV4t, pFST, and pMSTv) on the other hand could be homologous (Kolster et al., 2009; 2010). Yet, functional differences between both species have also been documented, since sensitivity to motion in area V3A and other regions within the intraparietal sulcus (IPS) was found to be more pronounced in human than in macaque (see Orban et al., 2003 for a review). For instance, cortical areas responsive to motion-defined structures are encountered in the human IPS but not in its monkey counterpart (Vanduffel et al., 2002).

Rather surprisingly, much less is known about the neural mechanisms that process motion along the depth dimension in primates. Motion in depth is nonetheless a very common and important component of motion in everyday life as it can notably signal objects moving towards the head and/or the body. Its estimation derives from two binocular cues: The change of disparity over time (CDOT), which tracks dynamic increase or decrease in horizontal disparity, and the interocular velocity difference (IOVD), based on opponent motion vectors between the two eyes (Nefs et al., 2010). The characterisation of the cortical areas processing each of these cues in both human and macaque would lead to a better understanding of how the 3D motion of objects is integrated in the primate brain and to further establish the similarities but also the differences between motion processing in the two species. Over the last ten years, this important line of research inspired a growing number of studies based on neuroimaging measurements in human and single-cell recordings in macaque.

In human, a pioneer fMRI study (Likova and Tyler, 2007) found that the strongest responses to CDOT arise in a cortical region anterior to the hMT+ complex. This region had reduced responses to 2D motion and was labelled ‘CSM’, for Cyclopean StereoMotion, by the authors. It might therefore be specialised in processing motion-in-depth. From additional analyses within independently defined regions of interest (ROIs), this study also found that several visual areas, such as V3A, V4, and hMT+, were more responsive to CDOT than to static disparity planes. Two years later, Rokers et al., (2009) found specific responses to CDOT and IOVD in the hMT+ complex, but also in area V3A and in lateral occipital regions (LO1/LO2). However, this study did not investigate activations in regions anterior to the hMT+ complex. More recently, Kaestner et al. (2019) confirmed that the hMT+ complex and two groups of ROIs that respectively gathered dorsal (V3A/B and IPS-0) and ventral (V4, LO-1 and LO-2) areas were more responsive to CDOT than to a control condition where the temporal frames were scrambled. These authors also found significant responses to CDOT in the CSM area but that were not as pronounced as those measured in hMT+. Altogether, these three human studies found consistent patterns of cortical responses to CDOT even though the precise functional role of area CSM, and notably how its responses to both 2D and 3D motion differ from those measured in hMT+, remains to be better understood.

In macaque, explorations of motion in depth selectivity at the single-cell level began with electrophysiological recordings in area MT, notably on the grounds that this area has similar 2D motion responses in human and non-human primates. Two studies found that MT neurons were selective to motion in depth (Czuba et al., 2014; Sanada et DeAngelis, 2014) but that this selectivity was primarily driven by the IOVD cue, with only a small contribution from the CDOT cue (Sanada & DeAngelis, 2014). Based on the human findings, it is possible that stronger responses to CDOT could be found in other satellite regions of the MT cluster (e.g. in FST or MSTv) and/or in more anterior regions of the STS. It is also possible that significant responses to CDOT exist in other regions of the ventral and dorsal visual pathways. In order to clarify which regions of the macaque cortex should be explored to better understand the neural mechanisms underlying motion-in-depth processing in primate, it is essential to first determine whether the areas that were identified in human from neuroimaging measurements could have putative homologues in macaques by using a similar approach.

In the present study, we performed fMRI recordings in awake behaving macaques to identify the cortical regions sensitive to motion-in-depth defined by changing disparity over time (CDOT). To facilitate the comparison with previous human data, we used an experimental protocol that was directly inspired from the neuroimaging studies described above (Likova and Tyler; 2007; Rokers et al., 2009; Kaestner et al., 2019). In order to precisely determine the limits of the MT cluster, we ran an additional retinotopic mapping experiment that permitted to delineate its four constituting areas V4t, MT, MSTv, and FST (Kolster et al., 2009). This approach allowed us to clarify whether the strongest responses to CDOT in macaque emerge beyond this cluster or within one or several of its components. It also permitted to document CDOT responses in retinotopic area of the early visual cortex (V1, V2, and V3) and along the dorsal (V3A) and ventral (V4) pathways. To determine whether regions activated by CDOT were also responsive to 2D motion, we ran a last experiment where we characterised the responses to moving versus static objects.

## Materials and Methods

### Subjects

Two female rhesus macaques (age: 15-17 years; weight: 5.4-6.2 kg) were involved in the study. Animal housing, handling, and all experimental protocols (surgery, behavioural training, and MRI recordings) followed the guidelines of the European Union legislation (2010/63/UE) and of the French Ministry of Agriculture (décret 2013–118). All projects were approved by a local ethics committee (CNREEA code: C2EA – 14) and received authorisation from the French Ministry of Research (MP/03/34/10/09). Details about the macaques’ surgical preparation and behavioural training are provided elsewhere (Cottereau et al., 2017).

### Data Availability

Data and analysis code will be made available after acceptance of the paper on a dedicated platform (OSF: https://osf.io/yxrsv/).

### Experimental design

Our stimuli were derived from those of previous fMRI studies that investigated how cyclopean stereomotion (motion-in-depth based on CDOT) is processed in humans (Likova & Tyler, 2007; Rokers et al.; 2009 and Kaestner et al., 2019). Our aim was to facilitate the comparison between the cortical networks involved in the two species. We used dynamic random-dot stereograms (dRDS) located within a disk (11 degrees of radius) and refreshed at 30Hz. The dot density was 15%. To manipulate binocular disparity between the two retinal projections, dots were green in one eye and red in the other and stimuli were observed through red-green anaglyph filters (stimulus code made available on OSF: https://osf.io/yxrsv/). In the ‘cyclopean stereomotion’ (‘CSM’) condition, dots within the upper and lower parts of the disk changed their disparity in opposition of phase, following a triangular function (1Hz) between ±23.3 arcmin (see figure 1-A). This disparity range was chosen to maximise the cortical responses to binocular disparities (see e.g. Backus et al., 2001; Cottereau et al., 2011). The opposition of phase between stereomotion of the dots in the upper and lower parts of the disc led to an average disparity across the visual field of zero at each frame and thereby prevented stimulus-induced change in vergence eye movement. Note that in this condition, motion in depth is defined from the change of disparity over time (CDOT). We chose to use the term cyclopean stereomotion in reference to the original human fMRI study of Likova and Tyler (2007).

**Figure 1:**
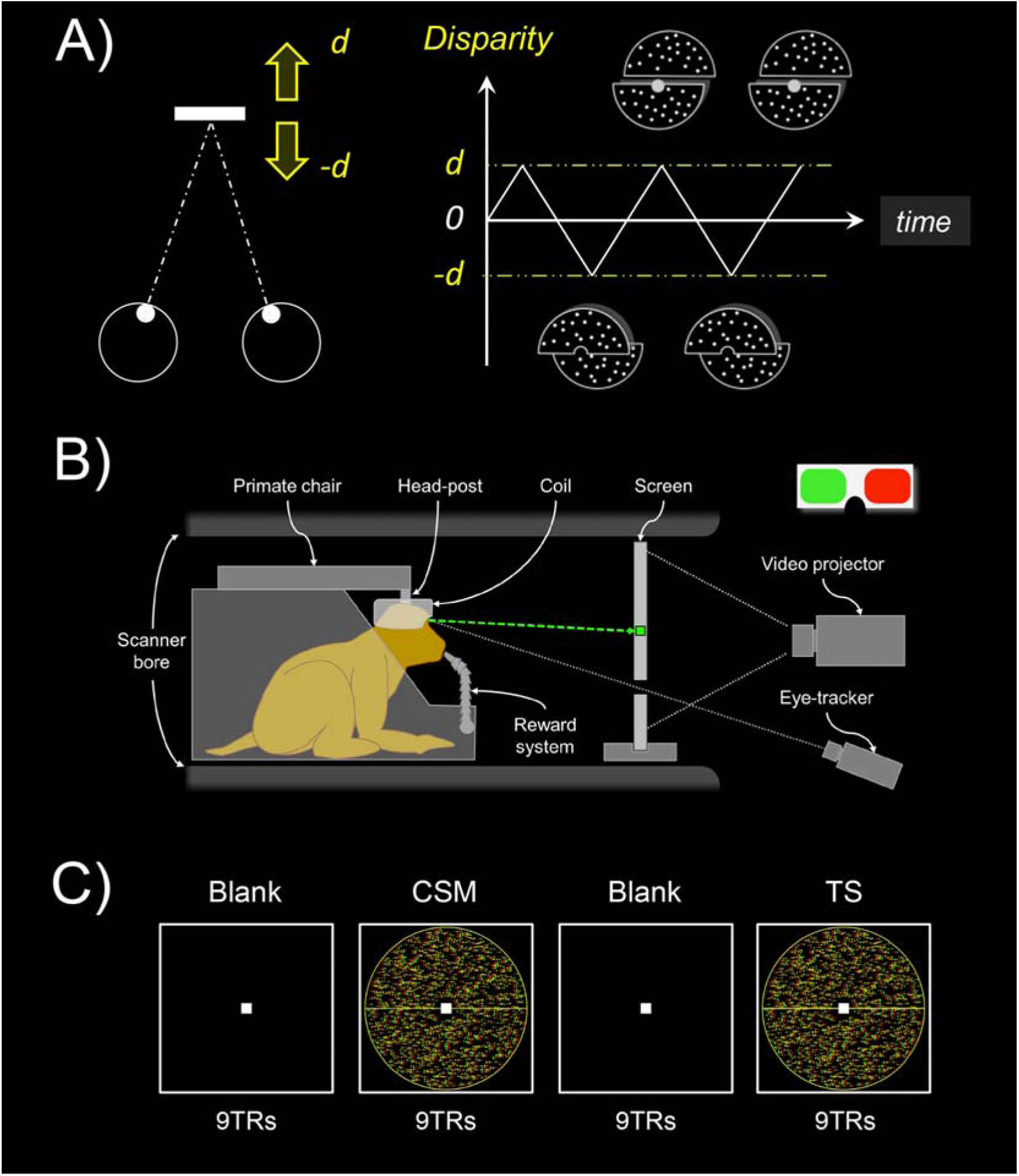
Stimulus design and experimental protocol. A) In the main condition (‘CSM’ for cyclopean stereomotion), the motion occurs along the antero-posterior axis (leftward panel). The stimulus consisted of a disk (11° of radius) defined by dynamic random dot stereograms (dRDS) refreshed at 30Hz. Its upper and lower parts moved in depth between d = ±23.3 arcmin in opposition of phase, following a 1Hz triangular function (rightward panel). In our control condition (‘TS’ for temporally scrambled), the individual frames of the CSM condition were shuffled in time to disrupt the smooth change of disparity over time. Our two conditions had identical retinal disparity distributions but only the CSM condition conveyed motion-in-depth. B) Schematic representation of the monkey fMRI setup. The animal was seated in a sphinx position within the primate chair, in the bore of the scanner, with the 8-channel, phased-array coil placed on top of the head. The animal was involved in a fixation task while its eye position was monitored by an infrared video-based eye-tracker. Horizontal disparity was introduced through red/green anaglyphs. C) Illustration of the experimental protocol. Recordings were performed using a blocked design, with the alternation of CSM and TS stimuli separated by blank periods. Each run contained 3 repetitions of such blocks plus an additional baseline period of 9TRs (117TRs in total). CSM conditions were shown first in half of the runs and TS conditions appeared first in the other half of the runs.

Our CSM stimulus led to perceive two planes continuously moving alongside a horizontal axis in opposite directions, one plane being located in front of the point of fixation and the other behind the fixation point. The control stimulus consisted of a ‘temporally scrambled’ version (‘TS’) of the CSM stimulus. To create this temporally scrambled condition, we shuffled the frames from the ‘CSM’ stimulus in order to disrupt the temporal sequence and thus, the motion in depth. Importantly, both conditions were monocularly identical and contained the same disparity distributions. The average relative disparities between dots in the upper versus lower parts of the disc were also identical between our two conditions.

### MRI recordings

#### Image acquisition: Templates of reference and functional sessions

Whole-brain images were acquired on a 3 Tesla MR scanner (Phillips Achieva) using a custom 8-channel phased-array coil (RapidBiomed) specifically designed to fit the skull of our macaques while preserving their field of view. Four T1-weighted anatomical volumes were acquired prior to the study for each monkey at a high resolution (MPRAGE; repetition time, TR = 10.3 ms; echo time, TE = 4.6 ms, flip angle = 8°; FOV: 155×155 mm; matrix size: 312×192 mm; voxel size = 0.5 x 0.5 x 0.5mm; 192 sagittal slices acquired in an interleaved order), as well as 300 functional volumes (gradient-echo EPI; TR = 2,000 ms, TE = 30 ms, flip angle = 75°, SENSE factor = 1.6; FOV: 100×100 mm; matrix size: 68×64 mm; voxel size = 1.25 × 1.25 × 1.5mm, 32 axial slices acquired in an interleaved order with a thickness of 1.5 mm and no gap). Those data were recorded in a single session whilst the macaques were slightly anaesthetised (Zoletil 100:10 mg/kg and Domitor: 0.04mg/kg) and their constants monitored with an MR compatible oximeter. Those volumes were then used to create individual anatomical and functional templates of reference.

Our T2*-weighted functional images were acquired with a gradient-echo EPI sequence with interleaved slice acquisition (TR = 2,000 ms, TE = 30 ms, flip angle = 75°, SENSE factor = 1.6; FOV: 100×100 mm; matrix size: 68×64 mm; voxel size = 1.25 × 1.25 × 1.5 mm, 32 axial slices acquired in an interleaved order with a thickness of 1.5 mm and no gap).

#### Scanning Procedure

During the functional recording sessions, macaques were head-fixed and seated in a sphinx position in a dedicated primate chair (see figure 1-B). They had to maintain their gaze within a central fixation window (2×2°) during daily sessions of up to 2 hours. Their fixation was monitored with an ASL© infrared video-based eye tracking setup at 60Hz and they were rewarded through a liquid delivery system (Crist Instrument) at intervals whose frequency depended on their fixation performance. Our cyclopean stereomotion stimuli were video-projected using a 23° x 23° field of view (viewing distance = 85cm). We used a blocked design based on cycles within which our two conditions (‘CSM’ and ‘TS’) were interleaved with baseline periods of fixation (see figure 1-C). Both the condition and baseline blocks lasted 18 seconds (9 TRs) and a cycle was therefore 72-second long (36 TRs). Each run contained 3 repetitions of this cycle plus an extra baseline that was added at the end for a total duration of 117 TR (234 seconds). We displayed the stimuli, controlled for the delivery of the liquid reward and the fixation performance using the EventIDE software (Okazolab®).

### Data analysis

#### Templates of reference

Anatomical and functional templates of reference were created for each individual with the volumes acquired prior to the current study. The anatomical template was obtained with the four T1-weighted anatomical volumes being realigned, averaged, and then co-registered on the MNI space of the 112RM-SL template (McLaren et al., 2009, 2010). To create the functional template, the 300 functional volumes (GE-EPI) were realigned, averaged, and then co-registered on the anatomical template. Both the T1 and the EPI mean images were segmented separately in order to obtain tissue probability maps for the grey matter, the white matter, and the cerebrospinal fluid (CSF). These probability maps were used to estimate the normalisation parameters from functional (mean EPI) to structural (mean T1) images for each individual.

#### Pre-processing of the raw functional data

Pre-processing and volume-based analyses were carried out with SPM12 in the Matlab environment (MathWorks®). Only runs with central gaze fixation above 85% were kept for further analysis. In total, we kept 43 and 60 runs for both macaques, respectively. The 4 first volumes of each run were discarded (dummy scans) to account for the establishment duration of the BOLD steady-state response. Pre-processing was performed for each volume, run by run. Slice-timing correction was performed first using as a reference the slice acquired in the middle of the acquisition of each TR. Images were then reoriented, co-registered with the EPI template, and transformed to fit the individual T1 template (with the normalisation parameters estimated between the mean EPI and T1 images; see the ‘*Templates of reference*’ section). No motion correction was applied to the images. Finally, the images were smoothed with a spatial Gaussian kernel (FWHM = 2×2×2 mm).

#### HRF estimation

Prior to our statistical analyses, we used independent datasets to characterise the BOLD haemodynamic impulse response functions (HRF) separately for each animal. These datasets respectively contained 16 (M01) and 12 (M02) 204s long runs that consisted of 6 cycles of 4s full field counter phasing (10Hz) checkerboards separated by a 30s blank interval (see more details about this procedure in Cottereau et al., 2017). Data were pre-processed using the pipeline described above and projected onto individual surfaces generated with the CARET software (Van Essen et al., 2001). Following Dumoulin and Wandell’s procedure (2008), we extracted the BOLD responses from nodes within the anatomically defined V1 of each individual. We only kept visually responsive nodes, that is those whose signal-to-noise ratio (SNR) was greater than 3. This SNR was estimated with a Fourier analysis of the average time courses across runs where the signal corresponded to the Fourier coefficient amplitude at the stimulation frequency F (i.e. F = 1/34) and the noise was given by the average moduli at the two neighbouring frequencies (i.e. F – δ f and F + δ f, where δ f = 1/2 is the resolution of our frequency analysis). We computed the average time course of these nodes during one cycle and used this average time course for estimating the HRF. The HRF was derived as the response to a 2s stimulus (our fMRI sampling rate). Note however that our stimulus duration was 4s rather than 2s because linearity deteriorates at short durations (Boynton et al. 1996; Logothetis and Wandell, 2004) and also because this duration was used in a previous monkey fMRI study that characterised the BOLD HRF in macaque (Leite et al., 2002). For each monkey, the average response to the 4s stimulus was fit as the convolution of the responses to two 2s responses, each of which is the HRF. We parameterised the HRF as the difference of two gamma functions (Friston et al., 1998). This functional form of the HRF captures the late undershoot of the response better than a single gamma function (Boynton et al., 1996).

#### General linear model (GLM) and whole-brain univariate analyses

Univariate statistics were performed at the voxel level in SPM12, using a general linear model (GLM). Our visual (CSM and TS) and baseline conditions were implemented as the three main regressors of the GLM. As reported above, we only analysed runs with fixation performance greater than 85%. We used the oculometric data of those runs to define regressors of non-interest that were included in the GLM to exclude the possible contribution of eye movements from our analyses. These regressors were obtained by automatically detecting the presence (1) or absence (0) of saccades in the different volumes of every run. The corresponding saccade regressors were then convolved with the HRF and introduced into the model. To characterise and eliminate noise in our recordings, we also performed a principal component analysis on voxels located outside the brain (see Farivar and Vanduffel, 2014). Indeed, time courses in those voxels should not be influenced by our experimental design but rather reflect artefacts caused by movements of the animal. For each run, we determined the number of principal components that were necessary to explain 80% of the variance in these voxels and used the corresponding principal vectors as regressors of non-interest in our model. This adaptive procedure typically added an average of 13.3 (±9.3) and 11.3 (±5.1) additional regressors in the models for Monkey 1 (MO1) and Monkey 2 (MO2), respectively.

We estimated the beta values associated with our GLM using the RobustWLS toolbox (Diedrichsen & Shadmehr, 2005), which is provided as an additional toolbox for SPM12 (http://www.diedrichsenlab.org/imaging/robustWLS.html). This approach allows estimating the noise variance for each image in the time series, using the derivative of a maximum likelihood algorithm. Variance parameters are then used to obtain a weighted least square estimate of the regression parameters of the GLM. It therefore helps to reduce the impact of noisier volumes on beta estimation. Previous studies showed that such a method significantly improved estimations in blocked-design fMRI experiments (see e.g. Takeuchi et al., 2011). The beta weights obtained from the GLM were subsequently used to perform univariate analyses (*t*-scores) at the whole brain level. These analyses were performed on the pre-processed EPI data and the beta weights were then projected onto the high-resolution volumes of our two animals. They were also projected on the individual cortical surfaces and on the cortical surface of the F99 template using the Caret software (Van Essen et al., 2001).

#### Localisation of areas selective to motion-in-depth and description of their responses

In order to identify areas with specific responses to motion-in-depth, we examined the statistical parametric map corresponding to the contrast between our two visual conditions (‘CSM’>‘TS’) and thresholded this map at *p*<10^−3^ (uncorrected, *t*-value>3.1). All the cortical regions that showed significantly stronger responses to CSM than to TS in both hemispheres and in each animal were considered. We controlled that these areas overlapped when projected on the right cortical surface of the F99 template. To further document the activations in those areas, we identified their local maxima and considered 3×3×3 voxel cubes around their coordinates. We then computed the percentage of signal change (PSC) corresponding to our main condition and its control using the following equations:

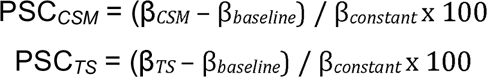

These values were extracted within small (3×3×3) voxel cubes rather than within patches determined by anatomical and/or statistical criteria, due to the fact that anatomical borders between areas are difficult to determine precisely and that our contrast led to extended activations that cannot be accurately divided into clusters corresponding to different functional regions (see figures 2, 3 and 4). Our approach is more conservative and avoids subjectivity when dealing with borders between areas. Importantly, we reproduced our analyses with betas extracted from smaller (1×1×1) or larger (5×5×5) voxel cubes, and this did not impact our results. Note that here we just document activations around the local maxima of selective areas (notably the relative difference between activations in our main condition and in its control and also the variability across runs) but we do not perform additional statistical analyses so as to avoid double dipping (Kriegeskorte et al., 2009).

**Figure 2:**
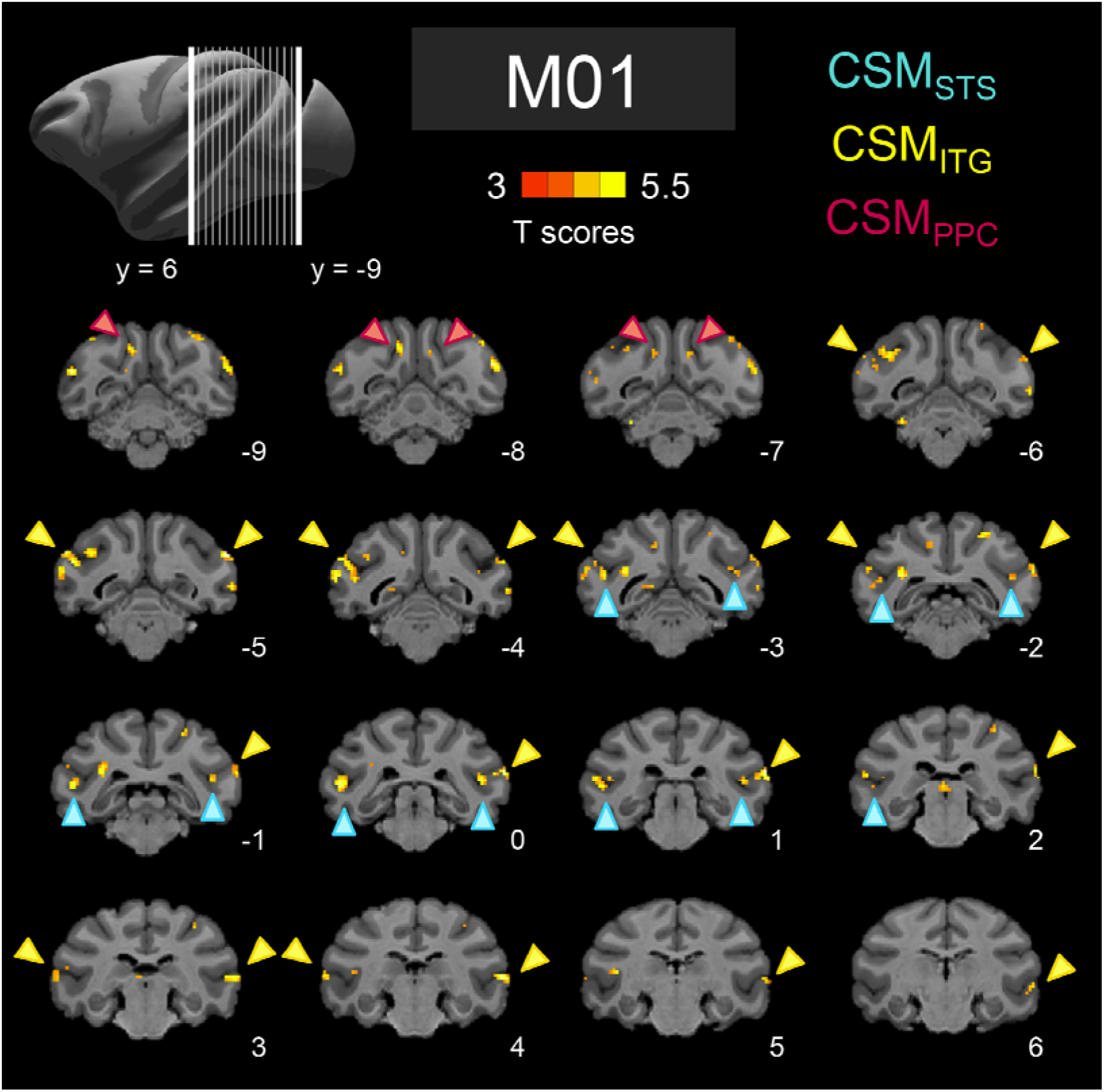
Activations for the contrast between Cyclopean Stereomotion (CSM) and its temporally scrambled version (TS) for M01. Figure shows activations that were stronger for the CSM condition than for the TS condition (T-score>3.1; p<10-3 uncorrected) Data are projected on the individual anatomical template of the macaque and are shown for different coronal slices. Coloured arrows indicate the localisation of our three regions of interest: CSM_STS_ (in blue), CSM_ITG_ (in yellow), and CSM_PPC_ (in pink). T-scores were obtained after computing the statistical parametric map for the contrast of interest between CSM and TS.

**Figure 3:**
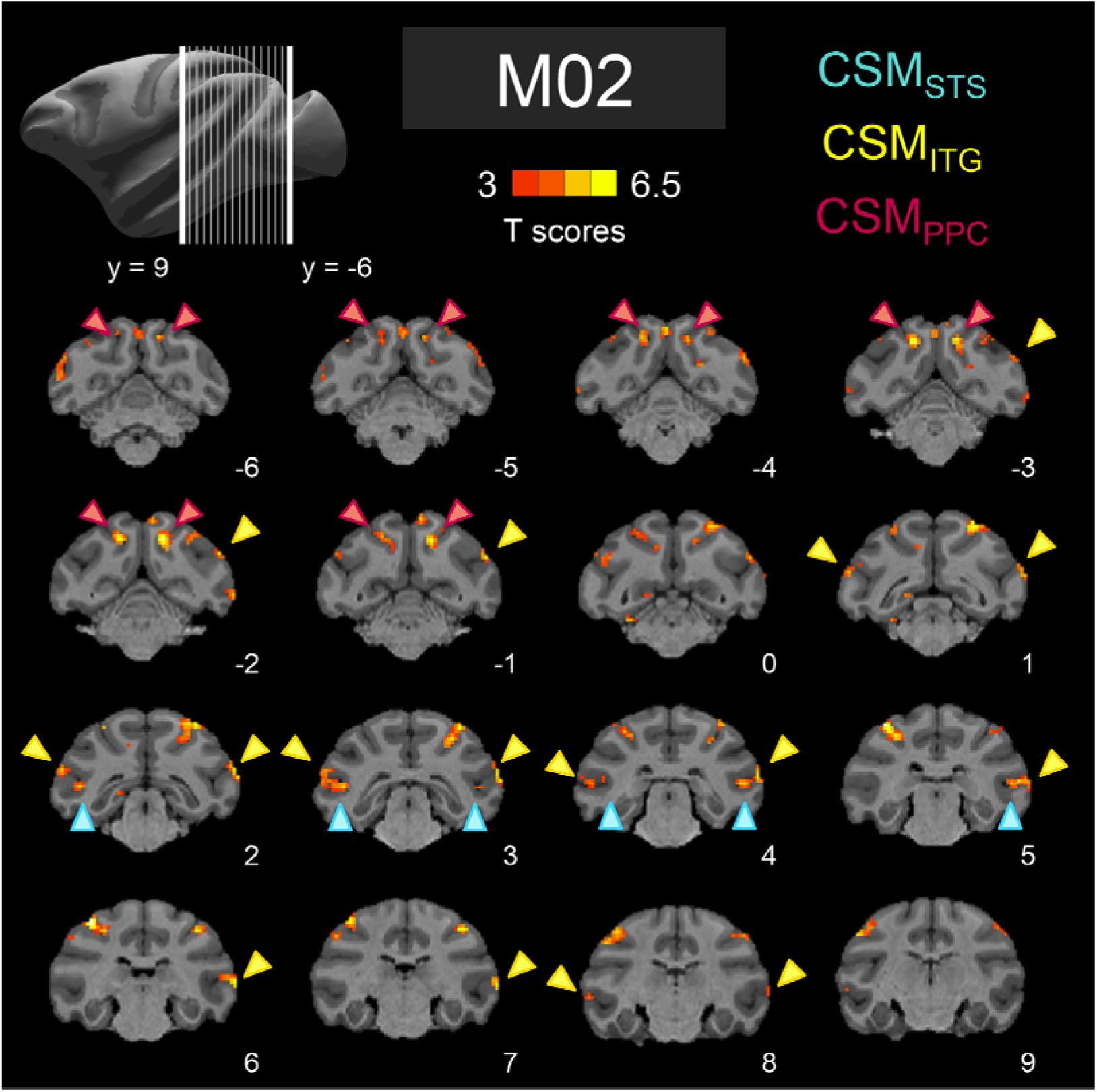
Activations for the contrast between Cyclopean Stereomotion (CSM) and its temporally scrambled version (TS) for M02. Conventions are similar to figure 2.

**Figure 4:**
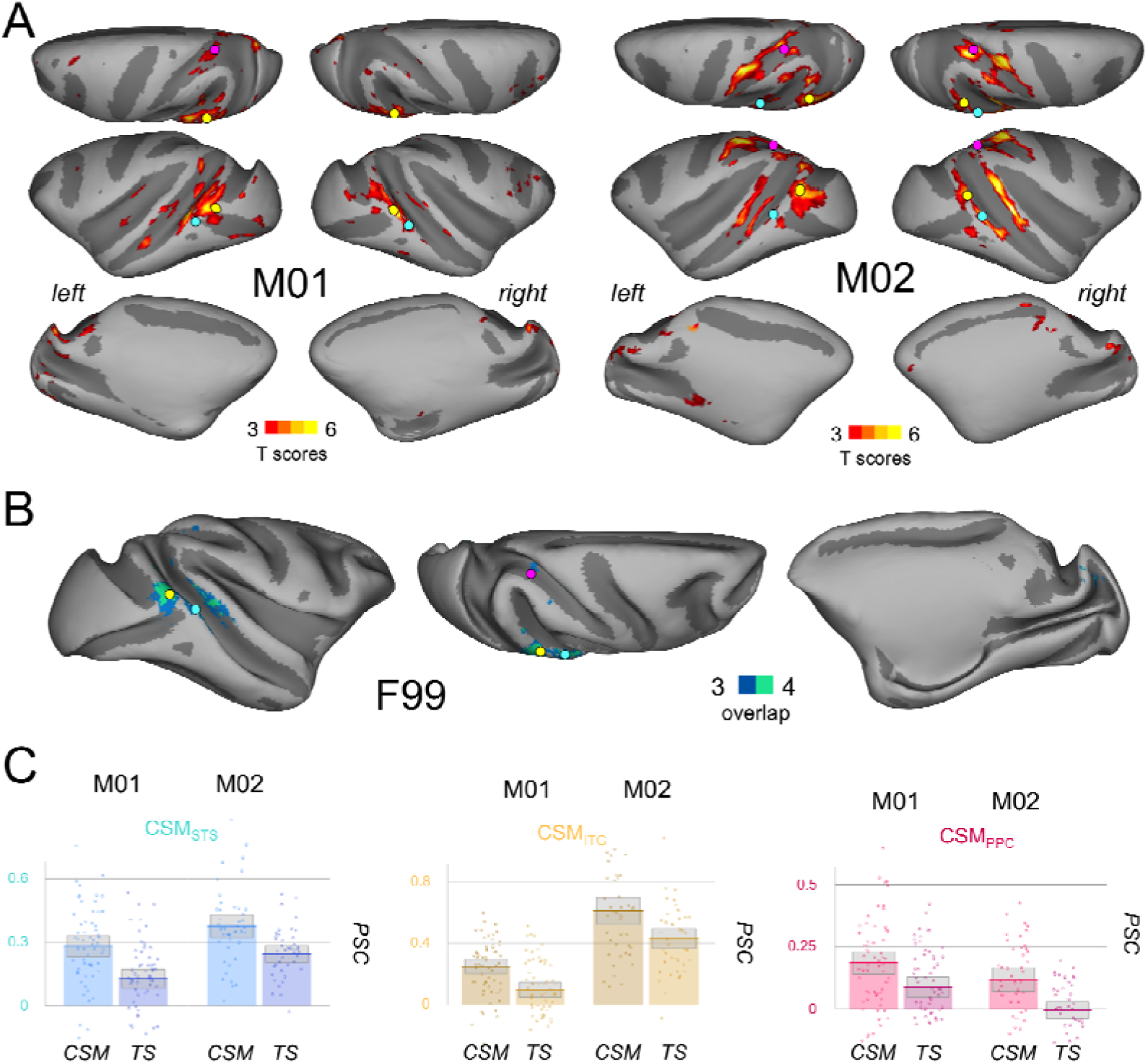
Activations for the contrast between Cyclopean Stereomotion (CSM) and its temporally scrambled version (TS) projected onto individual cortical surfaces and on the F99 template. A) Activations that were stronger for the CSM condition than for the TS condition. Data were thresholded at p<10^−3^ (uncorrected) and projected on the individual cortical surfaces of each animal (dorsal, frontal, and medial views). Coloured dots indicate the localisation of our three regions of interest: CSM_STS_ (in blue), CSM_ITG_ (in yellow), and CSM_PPC_ (in pink). B) Degree of overlap between the activations found in the two hemispheres of the two animals. The 4 individual cortical surfaces were morphed onto the right cortical surface of the F99 macaque template for projection of all the thresholded maps (frontal, dorsal and medial views). Blue colour indicates the overlap of 3/4 hemispheres and green colour of 4/4 hemispheres. C) Percentages of signal change (PSC) for the 2 visual conditions (CSM and TS) with respect to baseline (fixation on a black screen) in our three regions of interest. The boxes give the 95% confidence intervals for the average values. The dots provide the data for each run. A small jitter was introduced to facilitate visibility.

#### Definition of retinotopic areas and characterisation of their responses to motion-in-depth

We also performed a wide-field retinotopic mapping to delineate retinotopic regions that were used for additional ROI-based analyses. Whole-brain images were acquired with an identical setup as for the main experiment. In this case, visual stimuli were displayed using a large field-of-view (80° of visual angle, viewing distance = 25cm) and consisted of videos of a fruit basket that was moving laterally, forward and backward in monocular viewing. Traditional (clockwise/counter clockwise) rotating wedges (radius: 40°, angular extent: 49°) and expanding/contracting rings (eccentricity linearly varying between 0° and 40°) were used as visual apertures through which the fruit basket was displayed. Each run lasted 230s and contained 5 cycles of 44s with the first 10 seconds of a run being discarded (dummy scans) for the signal to reach its baseline. A small green square (0.4° x 0.4°) at the centre of the screen was used to control for fixation during passive viewing. As in our main experiment, only runs with more than 85% of correct fixation (respectively 47 and 48 runs for M01 and M02) were kept for further analyses. A pre-processing pipeline similar to the one described above was performed on the selected runs except that no smoothing was applied to the volumes and a fixed number of components (18 components) was used when performing the PCA, later used as a regressor of non-interest in the GLM. We projected the volume data onto individual surfaces using the Caret software (Van Essen et al., 2001) and a custom reorientation algorithm.

A population receptive field (pRF) analysis (Dumoulin & Wandell, 2008) was performed using the Matlab analyzePRF toolbox developed by Kay et al., (2013). For each surface node, an exhaustive set of theoretical pRF parameters (polar angle, eccentricity and size) was used to generate time courses that were compared to the real recordings. pRF size and position parameters that best predicted the data were selected as the node pRF parameters. With this approach, we obtained polar angle and eccentricity maps from which we characterised retinotopic areas that were described in previous monkey fMRI studies: V1, V2, V3, V4, V3A, as well as the regions within the Superior Temporal Sulcus (STS) (V4t, MT, MSTv, and FST) that form the MT cluster as described by Kolster et al. (2009). Those 8 retinotopically-defined regions were then projected back to the volumetric space to perform a ROI-based analysis of our motion-in-depth data. This was done using the inverse of the transformation between the volumetric and surface spaces mentioned above.

To test whether these retinotopic areas had specific responses to motion in depth, we first estimated their average PSC during the CSM condition and its TS control. We subsequently computed the corresponding difference between PSCs:

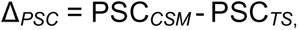

Note that we chose here to use the difference of PSCs because the PSCs for the CSM and TS conditions are paired. In order to estimate whether our observed PSC differences in these retinotopic areas were not due to chance, we computed permutation tests. We randomly attributed a negative sign to our PSC values and computed the mean difference, repeating this procedure 10,000 times. We then calculated a *p*-value defined as the proportion of random differences that were superior to our observed difference.

#### 2D motion localisers

To determine whether the regions that have specific responses to binocular 3D motion are also responsive to 2D motion, we performed a control experiment in which we contrasted responses to static images versus rich 2D motion stimuli. The scanning procedure was identical to the main experiment procedure. Motion localiser stimuli were based on the fruit basket video used for the retinotopic mapping experiment. For the static version, static images were randomly drawn from the video and refreshed at 1Hz. For the moving version, the video was normally played. Stimuli were displayed either centrally (<3° of eccentricity) or peripherally (>3° of eccentricity). As for the retinotopic experiment (see the previous section), these visual stimuli were displayed using a large field-of-view (80° of visual angle) at a viewing distance of 25cm. Each visual condition lasted 6 seconds and was interleaved with a 10-second baseline. The four visual conditions were presented in a pseudo-randomised order and were repeated 3 times within each run. Five extra baseline scans were added at the beginning of a trial for the signal to reach its baseline, thus resulting in a total duration of 202 seconds (101 TRs) for each run. In total 42 and 26 runs with fixation above 85% were kept for our analyses. Selected data was pre-processed as previously described, with an adaptive number of components that were necessary to explain 80% of the variance for each run, adding an average of 12.6 (±10.1) and 11.9 (±3.7) additional regressors in the model.

To estimate motion sensitivity in our regions of interest and in our retinotopic areas, we contrasted moving and static conditions, by combining central and peripheral presentations. We then performed a ROI-based analysis, looking at the BOLD activity within our independently defined regions.

## Results

The aim of this study was to identify the cortical network that processes disparity-defined motion-in-depth (i.e. cyclopean stereomotion) in two awake, behaving macaques using functional magnetic resonance imaging. Our experimental design was directly derived from previous human neuroimaging studies (Likova & Tyler, 2007; Rokers et al., 2009; Kaestner et al., 2019) so as to determine the homologies but also the differences between the BOLD activations in the two species (Orban, 2002). Our cyclopean stereomotion (‘CSM’) condition and its temporal scramble (‘TS’) control were defined from dynamic random dots stereograms (dRDS). They had identical retinal disparity distributions but differ in their temporal sequences (see the materials and methods section). Only the CSM condition conveyed motion-in-depth. Figures 2 and 3 show the statistical parametric maps obtained for the contrast between ‘CSM’ and ‘TS’ on the individual anatomical templates of each subject (M01 on figure 2 and MO2 on figure 3). These data are shown for different coronal slices.

On these two figures, red-to-orange colours indicate significantly stronger BOLD activations for the CSM condition than for the TS condition (*p*<10^−3^, uncorrected). Despite differences in the activation patterns observed in the two animals, this analysis reveals a network encompassing the temporal and parietal cortices in both monkeys. Notably, three cortical areas are consistently found in both the left and the right hemispheres of our two macaques. Coloured arrows show these areas. For sake of comparison with previous human neuroimaging studies, we named those areas after Likova and Tyler’s denomination (Likova & Tyler, 2007), that is, CSM for Cyclopean StereoMotion responsive areas. The first area (CSM_STS_) is located on the posterior bank of the superior temporal sulcus (STS) and extends posteriorly on the infero-temporal gyrus. The second one (CSM_ITG_) is located on the infero-temporal gyrus, at the intersection between the lunate sulcus, the inferior occipital sulcus (IOS) and the STS. The last area (CSM_PPC_) is localised in the posterior parietal cortex (PPC), mostly on the medial bank of the intra-parietal sulcus (even though activations can also be observed on its lateral bank in M02). This area might therefore correspond to the posterior intra-parietal (PIP) area. To be sure that the anatomical localisations of these 3 areas are not affected by our projections on the individual anatomical (T1) images, we confirmed their position on the functional (EPI) images in both monkeys (see supplementary figure 1). The MNI coordinates corresponding to the local maxima of these areas in the two animals are provided in table 1.

**Table 1:** MNI coordinates (in mm) of the local maxima for the 3 regions that were significantly more responsive for the CSM condition than for the TS control in the two hemispheres of the two animals.

| ROI | M01 |  |  | M02 |  |  |
| --- | --- | --- | --- | --- | --- | --- |
|  | x | y | z | x | y | z |
| **CSM<sub>STS</sub>** |  |  |  |  |  |  |
| **L** | -23 | 0 | 15 | -19 | 3 | 16 |
| **R** | 21 | 0 | 17 | 20 | 4 | 16 |
| **CSM<sub>ITG</sub>** |  |  |  |  |  |  |
| **L** | -29 | 4 | 15 | -23 | 3 | 18 |
| **R** | 28 | 1 | 20 | 26 | 3 | 19 |
| **CSM<sub>PPC</sub>** |  |  |  |  |  |  |
| **L** | -5 | -8 | 28 | -6 | -3 | 30 |
| **R** | 5 | -7 | 27 | 6 | -2 | 30 |

To demonstrate the consistency of these results across hemispheres, we show on figure 4 the projections of these activations on the individual cortical surfaces (see panel A).

As can be observed, our three regions of interest are found in all the individual surfaces, even though CSM_PPC_ is less visible in the right hemisphere of M01. This was confirmed by our projections of these activations on the right hemisphere of the F99 template. Figure 4-B shows that our three regions overlap in at least 3 hemispheres for CSM_PPC_ and in 4 hemispheres for CSM_STS_ and CSM_ITG_. The bar graphs on figure 4-C provide the activations elicited by our CSM condition and its TS control relative to baseline (blank screen) around those local maxima (see the material and methods). The thick lines provide the average values and the boxes give the corresponding 95% confidence intervals.

Finally, it is worth noting that in monkey M02, significant BOLD activations were also found in more anterior parts of the IPS notably within the ventral and anterior intraparietal areas (VIP and AIP, respectively). VIP has been shown to be involved in egomotion-compatible optic flow processing in both monkey (Cottereau et al., 2017) and human (Wall & Smith, 2008), whereas AIP has been suggested to play a role in 3D object processing and visually guided hand movements in both species as well (Sakata et al., 1997; Durand et al., 2007; Shikata et al., 2007). However, these activations were not confirmed in the second animal, potentially revealing a greater inter-subject variability in higher-order areas. In M02, activations were also found on the anterior part of the STS but they reflect responses from the fundus of the STS and/or from its posterior bank that were smoothed by our pre-processing pipeline and/or our transformations from the volume to the individual surfaces. The activations observed in the anterior part of the STS actually belong to clusters centred on the posterior bank.

Finally, it is important to emphasise here that we did not observe significant CSM responses on the cortex medial faces. Neither the anterior bank of the parieto-occipital sulcus, where the motion sensitive area V6 is located (see e.g. Pitzalis et al., 2013), nor the posterior part of the cingulate sulcus, where our group previously identified an area (pmCSv) responsive to egomotion-compatible optic flow (Cottereau et al., 2017), seem to have strong sensitivity to motion-in-depth.

### Retinotopic analysis

Previous studies in human found that the hMT+ complex had significant responses to stereomotion, notably based on changing disparity over time (CDOT) (Rokers et al., 2009; Joo et al., 2016). A single-cell study in macaque also found a weak but significant selectivity to CDOT in area MT (Sanada & DeAngelis, 2014). In order to determine whether the CSM-responsive ROIs we obtained from our univariate analyses overlap with (or correspond to) area MT and/or its neighbour regions, we performed a retinotopic mapping in our two animals (see more details in the Materials and methods section). This allowed us to delineate the areas of the MT cluster: V4t, MT, MSTv and FST (see Kolster et al., 2009), which is not possible with more classical motion localisers of the MT / hMT+ complex (even though some human studies proposed solutions to separate hMT from hMST, see Huk et al., 2002). In figures 5-A and 6-A, we show the locations of these areas and of our two CSM-responsive regions around the STS and the ITG, CSM_STS_ and CSM_ITG_, respectively. Figures 5-B and 6-B present the activations for the contrast between cyclopean stereomotion (CSM) and its temporally scrambled version (TS) projected on the same views.

**Figure 5:**
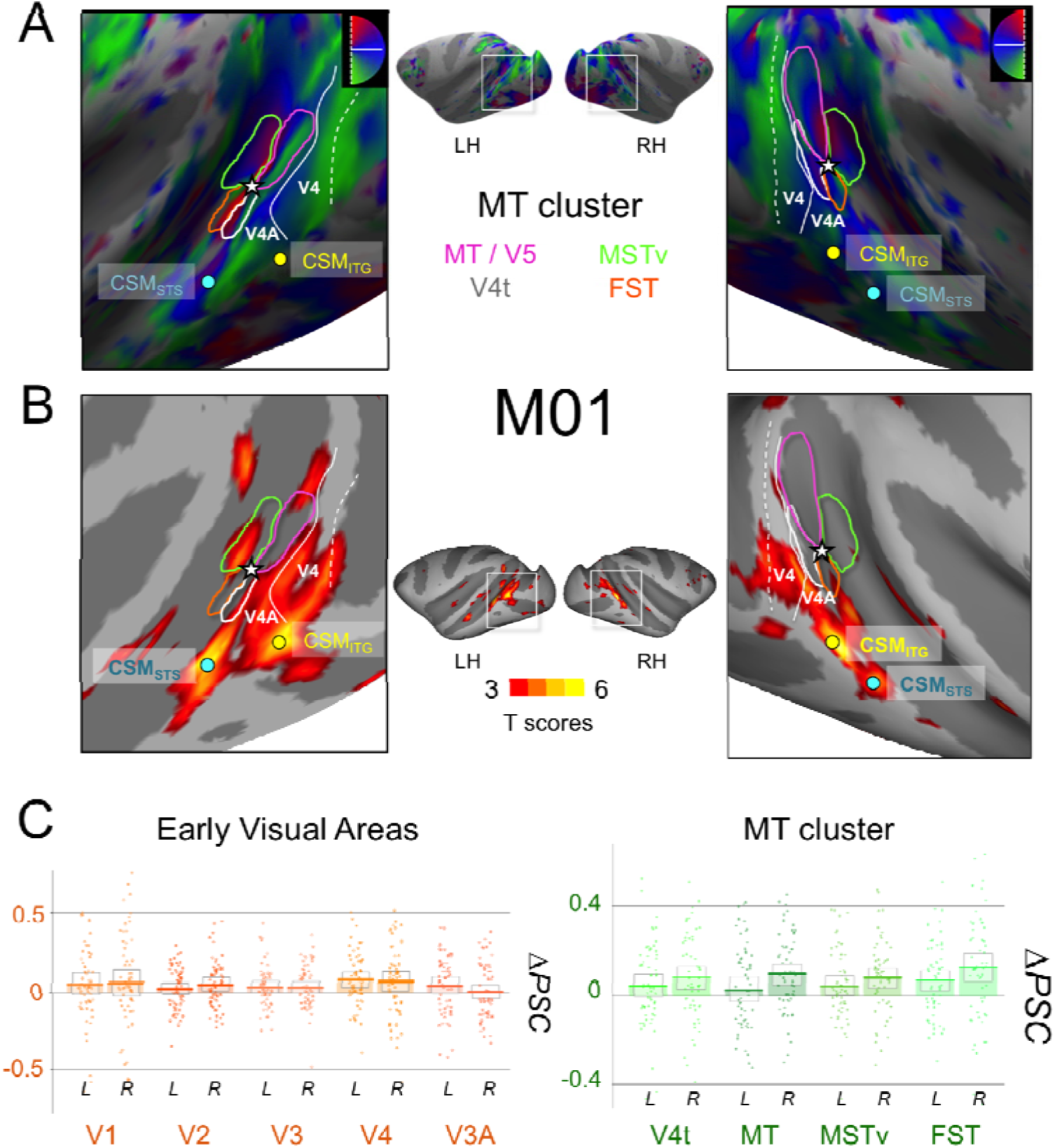
A) Retinotopic mapping of the Superior Temporal Sulcus (STS) for M01 and delimitation of the MT cluster areas: MT (dark blue), V4t (pink), MSTv (orange), and FST (green). We show here the polar angle maps that were used to delineate the borders between these areas. The extent of those areas was obtained from the eccentricity maps. We also show the representations of the vertical and horizontal meridians that delineate the borders between V3 and V4 and between V4 and V4A. Coloured dots indicate the local maxima positions for areas CSM_STS_ (in blue) and CSM_ITG_ (in yellow). As shown on the maps, CSM_STS_ and CSM_ITG_ are in the vicinity of the MT cluster, but clearly exterior to it. B) Activations for the contrast between Cyclopean Stereomotion (CSM) and its temporally scrambled version (TS) projected on the individual surfaces of M01, for both left and right hemispheres. C) Difference in percent signal change (ΔPSC) between the CSM and TS conditions in retinotopic areas. On the left, thick lines of the bar graphs provide average values for the left and right hemispheres of early visual areas: V1, V2, V3, V4, and V3A. On the right, average values are given for the MT cluster areas: V4t, MT, MSTv, and FST. The boxes give the 95% confidence intervals for the average values. The dots provide the data for each run. A small jitter was introduced to facilitate visibility.

**Figure 6:**
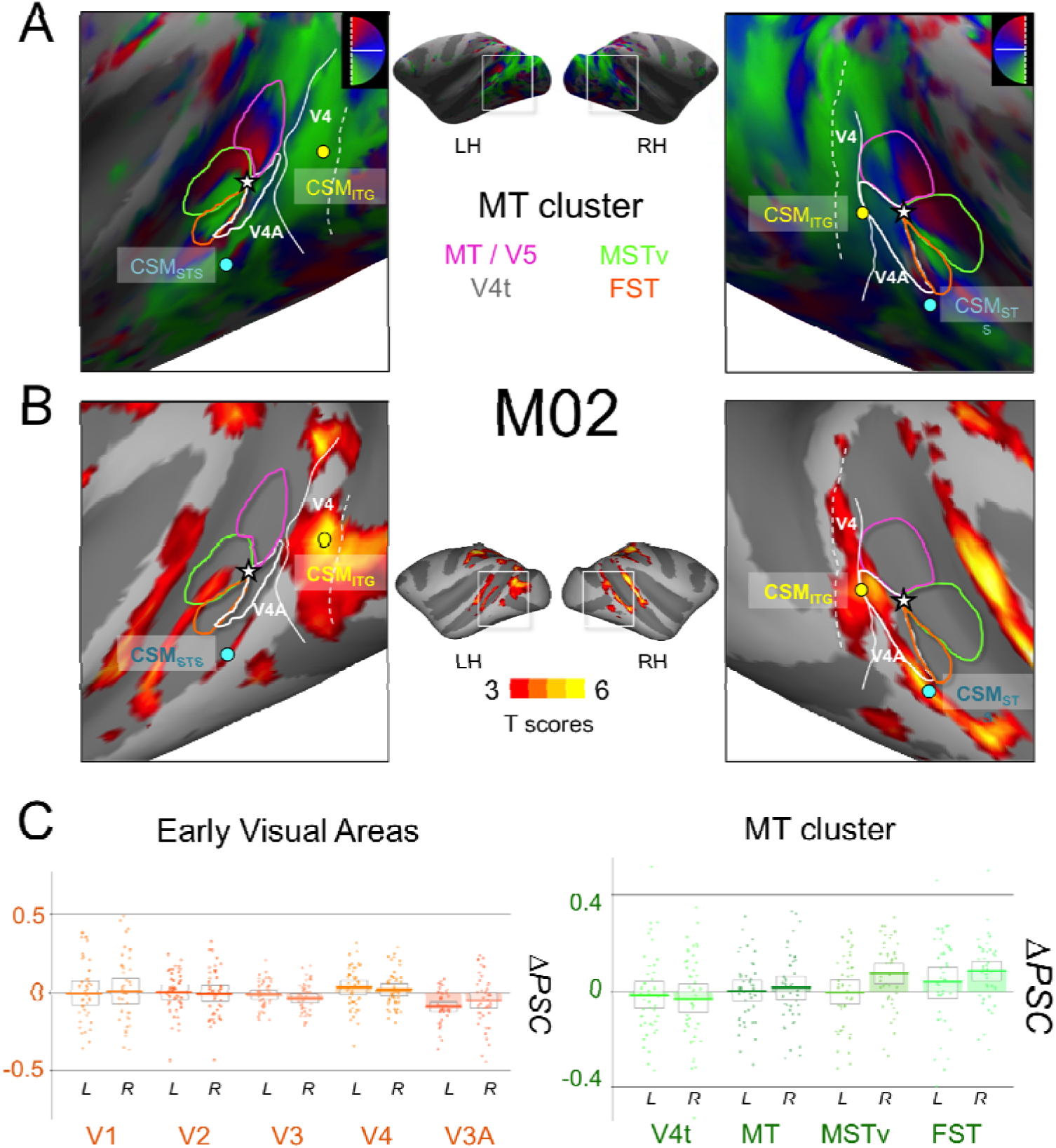
A) Retinotopic mapping of the Superior Temporal Sulcus (STS) for M02 and delimitation of the MT cluster areas: MT (dark blue), V4t (pink), MSTv (orange), and FST (green). B) Activations for the contrast between Cyclopean Stereomotion (CSM) and its temporally scrambled version (TS) projected on the individual surfaces of M02, for both left and right hemispheres. C) Difference in signal change (ΔPSC) between the CSM and TS conditions in retinotopic areas. Conventions are similar to figure 5.

We can see that although our two regions are close to the MT cluster, they do not overlap with it. Area CSM_STS_ is located more anteriorly along the posterior bank of the STS. Area CSM_ITG_ is located more posteriorly on the ITG, in a position that might correspond to areas V4 and/or V4A (see discussion).

To complete our study and to allow a direct comparison with previous human data (Likova and Tyler, 2007; Rokers et al., 2009; Kaestner et al., 2019), we performed ROI-based analyses within retinotopically-defined areas constituting the the MT cluster as well as the early-to-intermediate visual cortex: V1, V2, V3, V4, and V3A. The differences between the percentages of signal change (ΔPSC) for CSM versus TS in our two macaques are shown on figure 5-B and 6-B for early visual areas (V1, V2, V3, V3A and V4) and for the MT cluster (V4t, MT, MSTv, and FST). We can observe that if CSM selectivity in all these areas is not as pronounced as in CSM_STS_, CSM_ITG_, and CSM_PPC_, responses in the MT cluster tend to be stronger than those measured in V1, V2, and V3. Permutation tests demonstrated significant CSM effects in areas MT and V4t (1/4 hemispheres), MST (2/4 hemispheres, right hemispheres only), and FST (3/4 hemispheres). This suggests the presence of a marginal selectivity to cyclopean stereomotion in these regions. We also found that responses were significantly stronger for motion in depth in area V4 for one animal (2 hemispheres) but not for the other. Responses to CSM were not significantly stronger in V3A.

### 2D motion analysis

To test whether our three regions are only responsive to motion-in-depth or whether they are activated by motion in general, and notably by 2D motion, we ran an additional motion localiser in our two animals (n = 42 and n = 26 runs in M01 and M02, see more details in the Materials and methods section). We then computed the difference between the percentages of signal change (ΔPSC) corresponding to the 2D motion versus static image conditions. As expected from such a localiser, this analysis led to significantly stronger responses to motion in most of the retinotopic areas and more specifically within areas of the MT cluster. In particular, permutation tests demonstrated that all 4 regions of the MT cluster had significantly stronger responses to 2D motion in the two animals (*p*<0.05, except for left V4t in M02). We show in figure 7 the results of these analyses in our 3 CSM responsive areas (CSMSTS, CSMITG, and CSMPPC).

**Figure 7:**
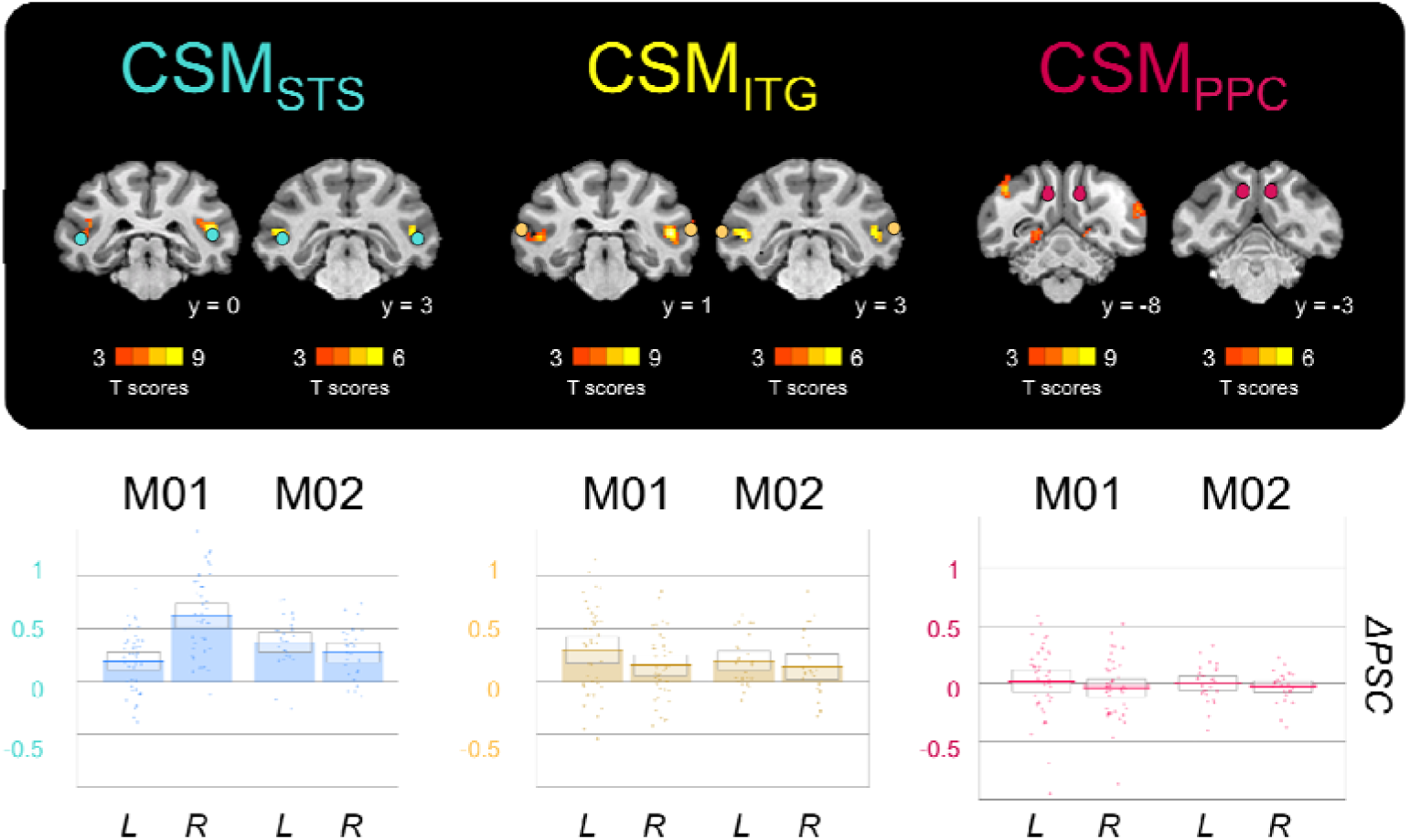
Sensitivity to 2D motion in CSM_STS_, CSM_ITG_, and CSM_PPC_. Stronger responses to 2D motion than to static snapshots of the same video sequences are shown on coronal slices from the individual anatomical template of each animal (upper panel). The colour dots provide the position of CSM_STS_, CSM_ITG_, and CSM_PPC_. For these 3 regions, PSC difference between responses evoked by 2D motion and static images is shown on the lower panel. The thick lines of the bar graphs provide the average values across runs for the left (L) and right (R) hemispheres of the two monkeys (M01 and M02). The boxes give the 95% confidence intervals for these average values. The dots provide the data for each run. A small jitter was introduced to facilitate visibility.

We can observe that only CSM_STS_ and CSM_ITG_ have a significant response to 2D motion (permutation tests, *p*<0.05), in both hemispheres for CSM_STS_ and in the left hemisphere for CSM_ITG_, for each monkey. Their motion selectivity (in particular in CSM_STS_) is therefore not specific to cyclopean stereomotion. On the opposite, responses to 2D motion in area CSM_PPC_ are not different from responses to static patterns (permutation tests, *p*>0.1). It implies that this region might uniquely respond to cyclopean stereomotion.

### Selectivity to 3D versus 2D motion within the lower bank of the STS

To further characterise the selectivity to 3D and 2D motion within the STS, we defined a path running along the posterior bank of this sulcus on the cortical surfaces of each hemisphere of our two animals. Each path departs from MT area and ends in the CSM_STS_ area. For each voxel along this path, we computed the average t-score within its first order neighbourhood (i.e. within a 3×3×3 cube centred on this voxel) for both the stereomotion versus temporal scramble and 2D motion versus static images contrasts. As t-scores for the second contrast were usually higher, we normalised the values along each path by dividing them by the maximum t-scores along the path. This facilitates comparisons between the sensitivity profiles for 3D and 2D motion. As shown in Figure 8-A, these profiles suggest that selectivity to stereomotion varies within the STS, with lower t-score values within the MT cluster and higher values in the CSMSTS area. By contrast, both the area CSM and the MT cluster exhibit strong sensitivity to 2D motion. This observation was confirmed by computing the average percentages of signal changes (ΔPSC) for 2D and 3D motion in those two regions (N.B.: the value for the MT cluster is the average ΔPSC across the path). The corresponding bar graphs are shown in Figure 8-B.

**Figure 8.**
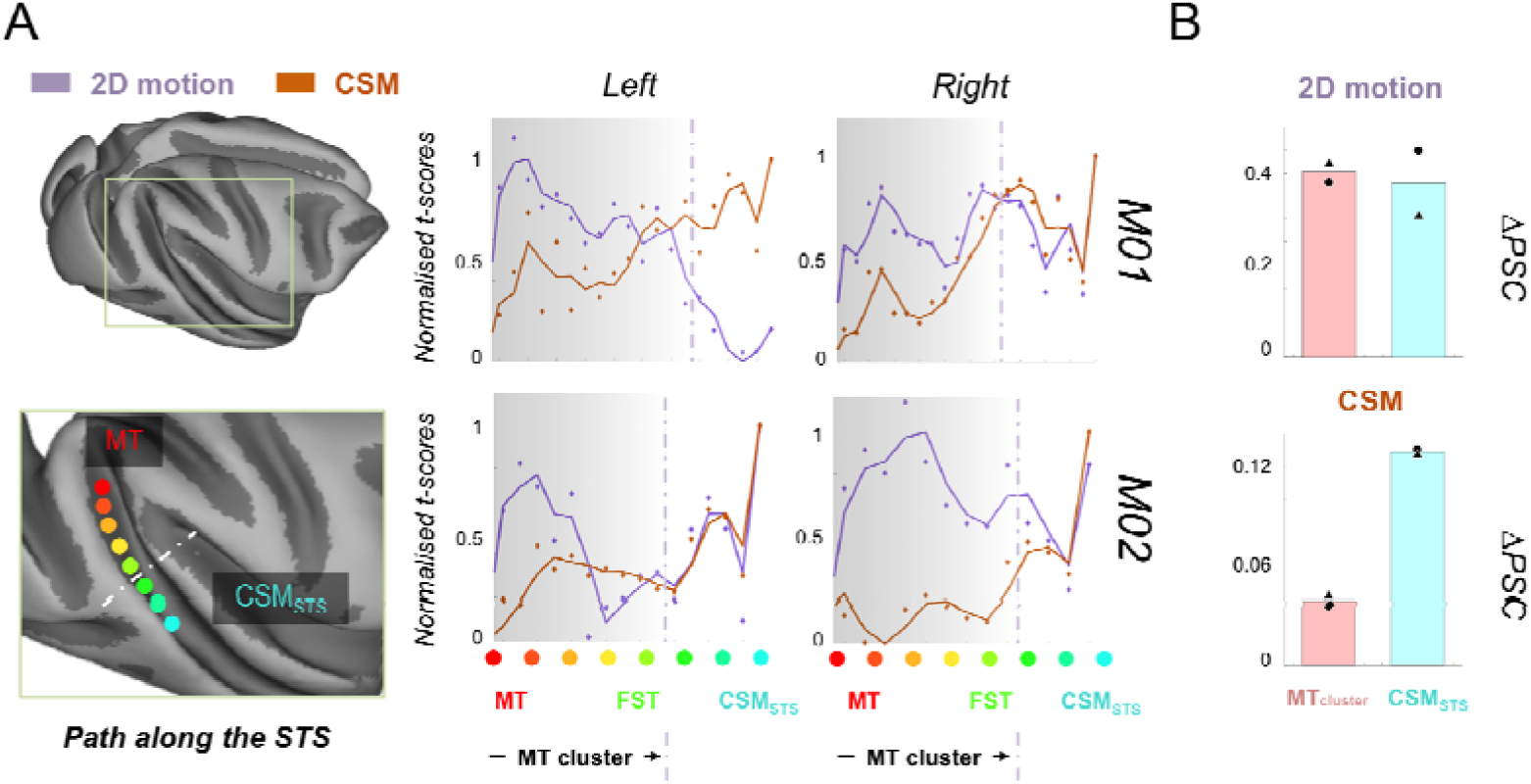
Selectivity to cyclopean stereomotion (CSM) and 2D motion within the STS. A) The left panel is a schematised view of the path drawn along the lower bank of the STS, starting from MT area (red dot) and ending within the CSM_STS_ area (cyan dot) defined from the stereomotion versus temporal scramble contrast. The grey dotted line represents the end of the MT cluster. On the right panel, responses to the cyclopean stereomotion (CSM, i.e. stereomotion versus its temporal scramble control) and 2D motion (i.e. 2D motion versus static images) contrasts are respectively shown in orange and purple for each hemisphere of both macaque subjects. Dots provide the normalised t-score values along the path whilst the curves were obtained from a median filtering of these values. The general trend is an increase of CSM selectivity along the STS, with the highest value outside the MT cluster. Selectivity to 2D motion peaks in the MT cluster and tends to decrease along the STS path. B) Average differences in signal change (ΔPSC) values in response to both types of tested motions (2D and CSM) in the four MT cluster areas (red bars) and in the CSMSTS area (cyan bars) for both subjects (triangle symbols for M01 and circle symbols for M02).

## Discussion

The aim of the present study was to characterise the cortical areas that process cyclopean stereomotion in rhesus macaque. To that end, we adapted the experimental protocols of previous human fMRI studies that investigated the cortical processing of motion in depth (Likova & Tyler, 2007; Rokers et al., 2009; Kaestner et al., 2019). Our main condition (‘CSM’ for cyclopean stereomotion) and its control (‘TS’ for ‘temporally scrambled’) shared the same disparity distribution and were monocularly identical (figure 1) but only the CSM condition conveyed stereomotion, since the temporal sequence was scrambled in the TS condition. We recorded whole-brain BOLD responses from two behaving macaques involved in a passive fixation task using a block design. Our analyses revealed a network of three areas whose responses to our CSM condition were consistently (i.e. across hemispheres and animals) stronger than those to our control condition (figures 2, 3, and 4). In reference to the original study of Likova & Tyler (2007), we labelled those regions CSM_STS_, CSM_ITG_, and CSM_PPC_. To complete these analyses, we also documented responses to our CSM condition in visual areas estimated from independent wide-field retinotopic mapping procedures (figures 5 and 6).

Our CSM_STS_ region is located on the inferior bank of the superior temporal sulcus (STS) and extends on the infero-temporal gyrus (see figures 2, 3, and 4), at a location (see table 1) anterior to the MT area and its satellites (V4t, MSTv, and FST, see figures 5 and 6). Based on our retinotopic maps, we confirmed that this region is outside the MT cluster (only a marginal overlap with area FST was found in the right hemisphere of M02). Our additional motion localiser demonstrated that it is also responsive to 2D motion (figure 7). Previous studies in macaque reported additional motion-sensitive regions in anterior portions of the STS. Using fMRI, Nelissen et al., (2006) notably documented an area in the lower superior temporal sulcus (LST) that responds to opponent motions and to actions. This area was 6-8mm anterior to FST and therefore does not fully coincide with our CSMSTS area. Another monkey fMRI study found several regions of the macaque inferior temporal cortex that had specific responses to disparity-defined stimuli (Verhoef, Bohon, and Conway, 2015). Among them, a region labelled ‘Pd’ (for posterior disparity) was localised in the lower bank of the STS, at a position that matches very well with our CSM_STS_ area. CSM_STS_ might thus be a distinct motion and disparity-selective area of the STS that notably processes motion in depth. In human, two studies (Likova and Tyler, 2007; Kaestner et al., 2019) found specific responses to cyclopean stereomotion in a cortical region anterior to the hMT+ cluster: CSM (N.B.: Rokers et al., (2009) only performed ROI-based analyses and it is therefore not possible to know whether they also had significant responses to motion in depth in this region). Our CSM_STS_ area might therefore be its macaque homologue. It is nonetheless important to note here that in the human studies, the delineation of the hMT+ complex (and sometimes its hMT and hMST sub-regions) was based on a contrast between the responses to uniform versus random motion whereas in our case the MT cluster was obtained from retinotopic mapping. To further clarify the potential homology between human CSM and macaque CSM_STS_, it would be interesting for future human studies to properly define area MT and its satellites using retinotopic mapping (see Kolster et al., 2010) in order to precisely determine the location of the CSM area with respect to those regions.

Our stereomotion contrast was based on a smooth variation in depth versus its temporally scrambled version (as in Kaestner et al., (2019) and in the ‘TS’ control of the second experiment of Rokers et al. (2009)). Although we used dynamic random dot stereograms, it is possible that this temporally scrambled control still evokes some apparent percept of motion. However, it lacked the smooth change of disparity of our main condition. In their experiments, Likova and Tyler (2007) used two planes that alternated between two different depths (thereby generating an apparent motion in depth) in their main condition whereas their corresponding control was a plane at a unique depth. We hypothesise that in both human and macaque, the CSM_STS_ / CSM area might be activated by different types of motion in depth, and notably by our contrast and the one used by Likova and Tyler (2007). This hypothesis is further supported by a control performed by these authors (see their supplementary materials) that demonstrated that significant activations were also obtained in this area with stimuli smoothly varying in depth (i.e. where binocular disparity was changed according to a sine wave), being therefore closer to those used in our own study.

Although our univariate statistics did not show significant responses in the MT area and its satellites, ROI-based analyses demonstrated that for some animal and/or hemispheres, responses in V4t (1/4 hemispheres), MT (1/4 hemisphere), MSTv (2/4 hemispheres), and FST (3/4 hemispheres) were significantly stronger for our stereomotion condition (figures 5 and 6). In a pioneer study, Maunsell and Van Essen (1983) concluded from single-cell recordings in area MT of anaesthetised macaques that neurons in this region had no selectivity to motion-in-depth (see also Felleman and Kaas, 1984). More recently, Sanada & DeAngelis (2014) found, using a different method, that MT does host neurons tuned to motion-in-depth (see also Czuba et al., 2014) but that these neurons were mostly driven by the inter-ocular velocity difference (IOVD) between the two eyes with only a modest contribution of the change of disparity over time (CDOT): ~10% of their neurons had significant selectivity for CDOT versus ~57% for IOVD. These findings are in line with our study and suggest that if selectivity to stereomotion is observable in area MT, it remains moderate. To our knowledge, selectivity to motion-in-depth was not directly tested in areas MSTv and FST. Based on our data, we hypothesise that a larger proportion of neurons tuned to cyclopean stereomotion could be found there. Altogether, the responses we measured in the STS are consistent with a model where stereomotion would be progressively integrated along a posterior-to-anterior axis with moderate responses in MT, intermediate responses in areas MSTv and FST, and stronger responses in CSM_STS_. This hypothesis is supported by our analysis of the responses on a path defined along the STS (see figure 8), which suggests that selectivity to stereomotion progresses beyond area MT.

Using a ROI-based analysis, all three previous human studies found significant responses to motion in depth in the hMT+ cluster. Likova and Tyler (2007) reported that selectivity in this cluster was weaker than in their CSM region, in agreement with what we found in macaque. In contrast, Kaestner et al. (2019) found that responses in hMT+ (in both hMT and hMST) were stronger than in CSM (see their figure 7). Given their use of a relatively small field of view (i.e. their stimuli had 5° of radius) contrasting with much larger stimuli in our experimental protocol (11° of radius) and in Likova and Tyler’s experiment (i.e. their display was a square of 30 x 40°), one possibility would be that neurons in the CSM_STS_ and CSM regions prefer more eccentric (>5°) stimuli. We computed the average population receptive field (pRF) eccentricities and sizes in the MT cluster and in CSM_STS_ (see supplementary figure 3-B and 3-D) and showed that in CSM_STS_ these parameters are actually similar to those found in V4t and FST, thus discarding this hypothesis. Further studies, notably in human where retinotopic mapping could be used to better define the position of CSM with respect to area MT and its satellites, will be needed to clarify this point.

Our CSMITG region is located on the infero-temporal gyrus, at the intersection of the lunate sulcus, the inferior occipital sulcus (IOS), and the STS (see figures 2, 3, and 4). It is therefore posterior to the MT cluster (see figures 5 and 6). This location matches well with area V4A that was previously described using single-cell recordings (Pigarev et al.; 2002) and fMRI (Kolster et al., 2014). CSM_ITG_ also overlaps with area V4, as suggested by our retinotopic analyses for which responses to motion in depth were significantly stronger in this area for one animal (see also panel B of figures 5 and 6). Unfortunately, signal-to-noise ratios in our retinotopic data were not sufficient to properly map area V4A (notably its anterior border with area OTd) and further studies will be needed to clarify its responses to stereomotion relatively to those estimated in V4. Both V4 and V4A are activated by disparity (Watanabe et al., 2002; Verhoef et al., 2015) and motion (Li et al., 2013; Kolster et al., 2014), even though their motion selectivity is not as pronounced as in the MT cluster (Kolster et al., 2014). This in line with our finding that CSM_ITG_ has only moderate responses to 2D motion, notably when compared to motion responses in area CSMSTS (figure 7). Interestingly, two human studies on motion in depth (Likova and Tyler, 2007; Kaestner et al., 2019) reported significant CSM responses in area V4. Kaestner et al. (2019) also found strong steremotion responses in area LO-1 (and Rokers et al. (2009) in area LO which includes LO-1) which was proposed to be the human homologue of area V4A (Kolster et al., 2014). These results suggest a good correspondence between CSM responses in those regions of the human and macaque brains.

Area CSM_PPC_ is localised in the posterior parietal cortex (PPC), mostly on the medial bank of the intra-parietal sulcus (IPS). Responses in this region were not stronger for 2D motion than for static stimuli, in agreement with previous monkey fMRI studies (see e.g. Vanduffel et al., 2001). Because of its localisation, area CSM_PPC_ might correspond to the posterior intra-parietal area (PIP) (Colby et al., 1988; Markov et al., 2014). Even though further studies will be needed to clarify this point, it is tempting to hypothesise that there might be a functional dissociation for 3D processing between this area and its counterpart on the lateral bank of the IPS, the caudal intra-parietal area (CIP). Indeed, in a previous monkey fMRI study, Durand et al. (2007) revealed sensitivity to kinetic depth in area PIP and AIP (for which we also found activations in M02) but not in area CIP. Area PIP has also been shown to respond to 3D structure (see e.g. Alizadeh et al., 2018). It might therefore play a role in the detection of and interaction with moving objects whereas CIP could be involved in processing 3D orientation and/or 3D features/arrangement of elements (Tsutsui et al, 2002; Durand et al., 2007; Rosenberg et al., 2013). In human, the studies of Likova & Tyler (2007) and of Rokers et al. (2009) did not explore stereomotion selectivity in the parietal cortex (the latter nonetheless reported significant responses to CDOT in dorsal area V3A). The only study that reported results at the whole-brain level (Kaestner et al., 2019) found strong stereomotion responses in area IPS-0, which is located in the caudal part of the human IPS and therefore constitutes a potential homologue of our CSM_PPC_ region. Further studies will be necessary to clarify this point.

Finally, we did not find CSM specific activations in area V3A although this area is motion selective in macaque (Galetti et al., 1990). Stereomotion significantly activated human area V3A in previous studies (Likova & Tyler, 2007; Rokers et al., 2009; Kaestner et al., 2019). In this area, this is not the only functional difference in motion processing between the two species as responses to structure from motion are known to be much stronger in human V3A than in macaque V3A (Vanduffel et al., 2002, see also Orban et al., 2003).

In order to avoid eye movements, and notably vergence, to contaminate our activations, we took several precautions. As mentioned in the Material and methods section, we only kept runs for which fixation performance was above 85%. Furthermore, our stimuli were designed to avoid driving vergence, with an average disparity value across space that was always equal to zero. Finally, we used the detected saccades as regressors of non-interest in our GLM. It is also worth noting that the activations we observed when contrasting our two conditions of interest (CSM vs. TS) are different from the vergence networks as investigated in terms of vergence tracking and vergence steps by Ward and collaborators (Ward et al. 2015). Additional analyses based on fixation performances and variance of eye position along the x and y axes during the CSM and TS conditions further demonstrated that eye movements did not impact our results (see supplementary figure 2 and the accompanying text).

## Conclusion

Our fMRI recordings in two macaques demonstrated that cyclopean stereomotion is mainly processed by three cortical areas: CSM_STS_, CSM_ITG_, and CSM_PPC_. We also observed a moderate selectivity in areas of the MT cluster, mostly MSTv, and FST. These results are close to those described in human using a similar experimental protocol and therefore suggest that the cortical network processing stereomotion is relatively well preserved between the two primate species.

## Acknowledgments

This work was supported by the Agence Nationale de la Recherche Jeunes Chercheuses et Jeunes Chercheurs (Grants ANR-16-CE37-0002-01, 3D3M) and an ATP funding from the Paul Sabatier University awarded to B.R.C. We thank the Inserm/UPS UMR1214 Technical Platform for the MRI acquisitions.

