## Supplementary figures 1,2,3 for "Stereomotion processing in the non-human primate brain"

The supplementary materials contain 4 items: one supplementary text (‘Influence on eye movements’) and 3 supplementary figures.

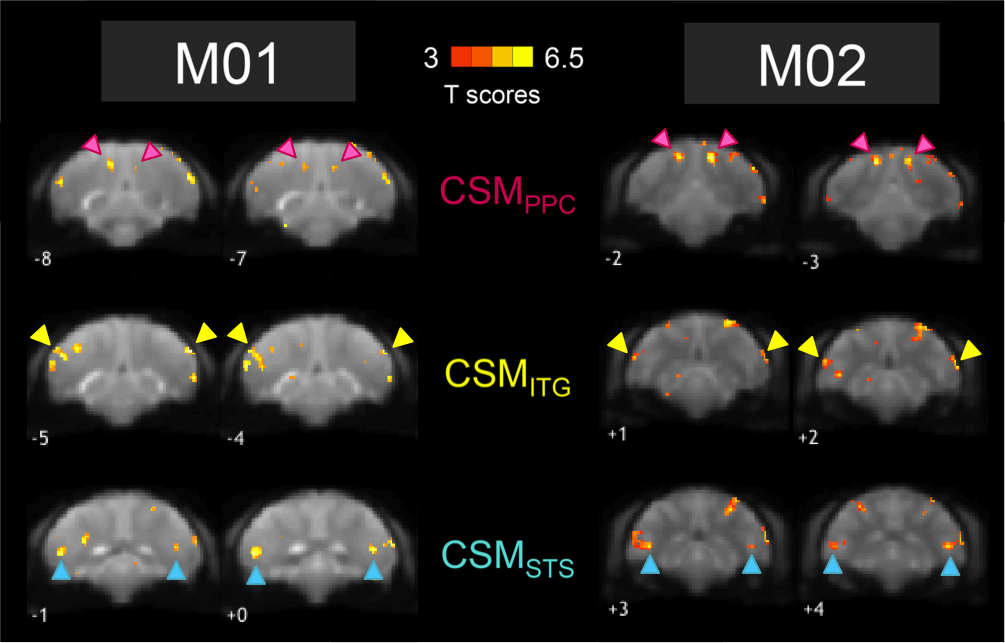
***Supplementary Figure 1:*** *Activations for the contrast between Cyclopean Stereomotion (CSM) and its temporally scrambled version (TS) for M01 and M02. Figure shows activations that were stronger for the CSM condition than for the TS condition. Data are projected on an individual average EPI image of each of the macaque and are shown for different coronal slices. Coloured arrows indicate the localisation of our three regions of interest: CSM_STS_ (in blue), CSM_ITG_ (in yellow), and CSM_PPC_ (in pink). T-scores were obtained after computing the statistical parametric map for the contrast of interest between CSM and TS.*

%%%%%%%%%%%%%%%%%%%%%%%%%%%%%%%%%%%%%%%%%%

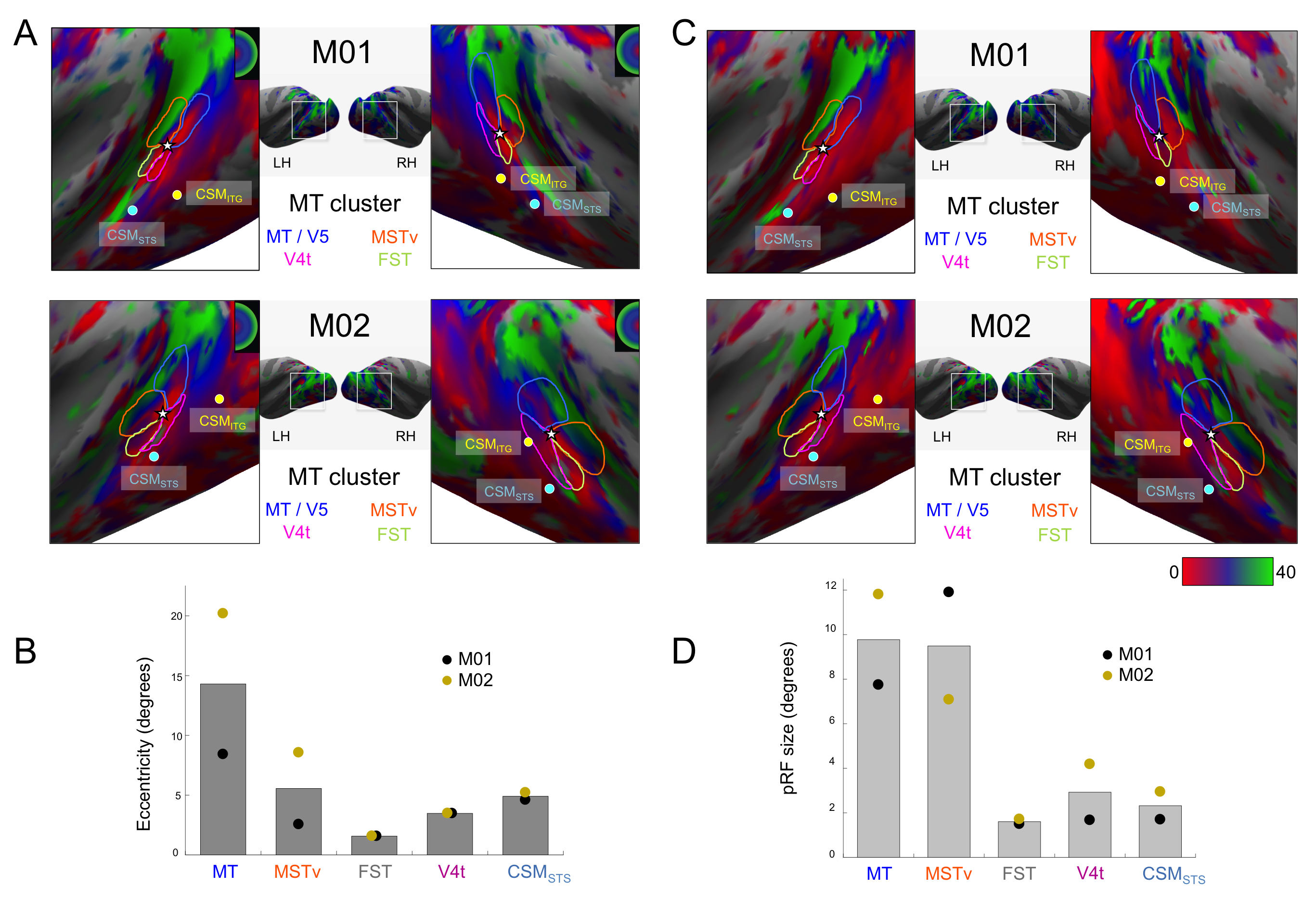

***Supplementary figure* *2*:** *A) Retinotopic mapping of the Superior Temporal Sulcus (STS) and delimitation of the MT cluster areas: MT (dark blue), V4t (pink), MSTv (orange), and FST (green). We show here the eccentricity maps of both subjects (M01 and M02) and the local maxima positions for areas CSM_STS_ (blue dot) and CSM_ITG_ (yellow dot). B) Average eccentricities for the four areas of the MT cluster (same colours as in A) and for CSM_STS_ for both subjects (M01 in black and M02 in gold). C) Retinotopic mapping of the Superior Temporal Sulcus (STS) for M01 and delimitation of the MT cluster areas: MT (dark blue), V4t (pink), MSTv (orange), and FST (green). We show here a map of the population receptive fields (pRF) size for both subjects (M01 and M02) and the local maxima positions for areas CSM_STS_ (blue dot) and CSM_ITG_ (yellow dot). D) Average pRF sizes for the four areas of the MT cluster (same colours as in C) and for CSM_STS_ for both subjects (M01 in black and M02 in gold).*

%%%%%%%%%%%%%%%%%%%%%%%%%%%%%%%%%%%%%%%%%%

***Influence of eye movements***

To confirm that eye movement did not affect our results, we further analysed the oculometric data, by determining whether fixation performances significantly varied between our two conditions of interest (CSM and TS). The distributions of these performances in our two animals for the two conditions are shown in supplementary figure 2. Given that these data were not normally distributed, this was done using permutation tests (see the Materials and Methods for more details). We did not find any significant difference in fixation performances between our two conditions in M02. In M01, fixation scores were slightly higher for the TS condition (*p* = 0.0124). Nonetheless, these differences were not correlated with differences in PSC in our three ROIs (CSM_PPC_, CSM_ITG_, and CSM_STS_). We therefore conclude that our results were not driven by differences of fixations scores between our two conditions. We use the same approach to analyse the variance in eye fixation along the x and y-axes. Variances in X and Y coordinates did not vary between our two conditions in M01. In M02, variance in X and Y were slightly higher for the TS condition but here as well, we did not find correlation between these differences in variance and difference of PSC in our 3 ROIs. It is thus highly unlikely that the ocular behaviour of our subjects had any impact on our results.

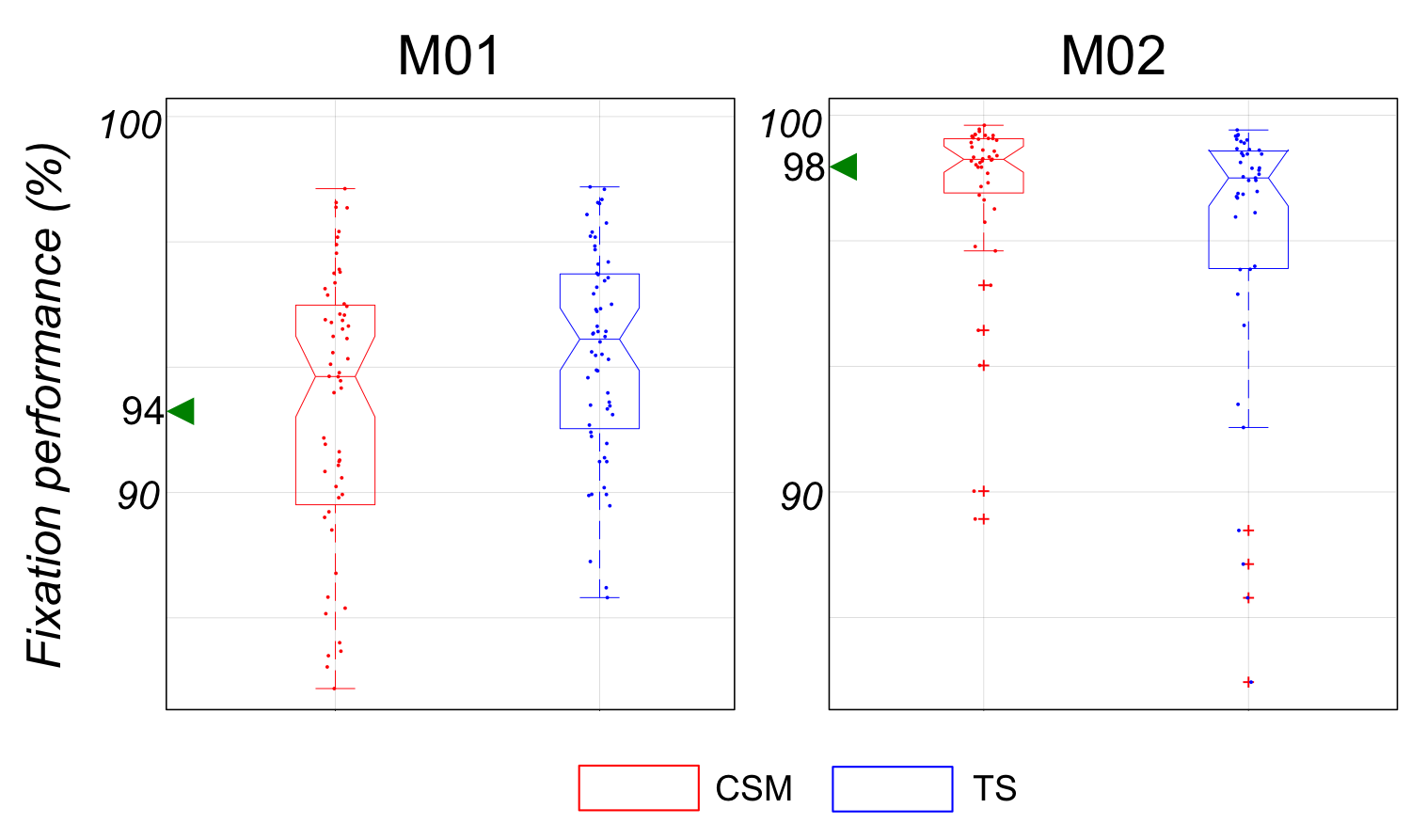

***Supplementary figure 3:*** *Ocular behaviour. Boxplots represent the distribution of percentages of correct fixation (i.e. less than 2° away from the fixation point) for both CSM and TS conditions and for each monkey (median and interquartile values). Numbers and arrows in green indicate the median values of the percentage of fixation performance during the baseline condition. Jittered dots are the fixation performance values for every run.*

%%%%%%%%%%%%%%%%%%%%%%%%%%%%%%%%%%%%%%%%%%
